# Stroke and Alzheimer’s disease have distinct consequences on neurovascular function but synergize to increase amyloid deposition

**DOI:** 10.64898/2026.08.29.746901

**Authors:** Ozama Ismail, Simone S. Woodruff, Benjamin Zimmerman, Wenri Zhang, Teresa L. Stackhouse, Martin M. Pike, Randall L. Woltjer, Anusha Mishra

## Abstract

**INTRODUCTION:** We examined whether stroke induces chronic cerebrovascular dysfunction and thereby exacerbates Aβ pathology.

**METHODS:** We exposed Tg2576 mice to a transient mild subcortical ischemia and examined cerebrovascular function at chronic timepoints as well as reactive astrocytes and Aβ deposition post-mortem.

**RESULTS:** Baseline cerebral blood flow and cerebrovascular reactivity (CVR) are both influenced by anesthesia regimen. Aβ strongly impairs CVR at earlier ages, while mild ischemia impairs CVR at later ages. Mild ischemia reduces neurovascular coupling more strongly than Aβ and synergistically increases Aβ deposition in Tg2576 mice.

**DISCUSSION:** Stroke-induced cerebrovascular dysfunction and astrocyte reactivity persist for long periods after the injury. Although there is variability in the spatiotemporal progression of ischemia-and Aβ-induced vascular impairments, they generally follow the pattern of Thal staging. Surprisingly, the ischemia+Aβ group showed the least cerebrovascular dysfunction yet an exacerbation of Aβ deposition, suggesting that these pathologies are connected but not tightly coupled.

## 1 INTRODUCTION

As the rates of Alzheimer’s disease (AD) climb globally^1^, it is fast becoming one of the greatest healthcare challenges of our time. Despite significant advances in the field, the precise mechanisms behind disease onset remain poorly understood, especially for sporadic AD, and represent a significant hurdle in the race to find a cure. Although classically characterized by the aberrant accumulation of amyloid beta (Aβ) plaques^2^ and tangles of phosphorylated tau^3^, recent cohort studies indicate that cerebrovascular dysfunction in the AD brain are one of the earliest measurable changes that precede dementia-associated decline^4^. That this relationship emerged even in the AD Neuroimaging Initiative dataset, especially in the form of white matter hyperintensities and silent infarcts, despite the inclusion criteria selecting against vascular risk factors^5^ is a telling indication of the close relationship between vascular disease and AD.

Ischemic injuries are of particular interest in AD, given that those who suffer a stroke are at higher risk of developing long term cognitive decline^6^. Further, imaging studies have revealed that aging patients who harbor silent infarcts caused by small, asymptomatic strokes are at double the risk of dementia^7, 8^. Silent strokes are present in almost 35% of people above 85 years of age^9^ and microinfarcts are observed in AD brains up to 10 times more frequently than healthy controls^10^. Furthermore, post-mortem findings indicate that AD patients with infarcts experience more severe cognitive decline than those with AD pathology alone^11^. Given that aging and stroke are both strong risk factors for AD and related dementias (ADRD)^12^ and the earliest pathophysiological changes in the ADRD brain are of vascular origin^4^, it is possible that stroke induced vascular dysfunction is an early event in ADRD. Indeed, in patients with stroke, cerebrovascular reactivity (CVR) and neurovascular coupling (NVC) are both attenuated in regions beyond the infarct^13–15^, and cognitive dysfunction observed in stroke patients is attributed to this consequence^16^. We previously demonstrated that experimental stroke in animal models also similarly diminishes NVC responses in regions outside infarct ^17^, and others have shown that, in rodent models, stroke increases abnormal processing of amyloid precursor protein by 30%^18^. This suggests that small strokes could contribute to ADRD pathology in mixed dementia through chronic cerebrovascular dysfunction.

To understand this etiology, we developed an innovative experimental model of mixed dementia by exposing Tg2576 mice, a transgenic model overexpressing amyloid beta (Aβ), and their wild-type (WT) littermates to a model of transient mild subcortical ischemia (tMSCI). We then characterized changes in CVR and NVC at chronic timepoints after the ischemic insult and assessed reactive astrocytes and Aβ deposition post-mortem. Our results reveal that Aβ strongly contributes to CVR impairments at earlier ages, while ischemia impairs CVR at later ages. Regional analysis revealed that the cortex, hippocampus, and amygdala were more vulnerable to these changes than the thalamus. Further, we found that ischemia has a stronger impact on lowering NVC than Aβ. We also observed a synergistic effect on Aβ deposition, as evidenced by larger deposit sizes in Tg2576 mice exposed to mild ischemia.

## 2 METHODS

### 2.1 Animals

All experiments were approved by the Oregon Health & Science University Institutional Animal Care and Use Committee. This study involved the use of Tg2576 mice and their WT littermates up to 15 months (mo) old (**Figure 1A**). The Tg2576 mice were obtained via internal transfer from Dr. Jeffrey Iliff (previously at Oregon Health & Science University, currently at University of Washington). Mice were continuously bred under our care, and genotyping was performed through Transnetyx (Memphis, TN, USA).

**Figure 1.**
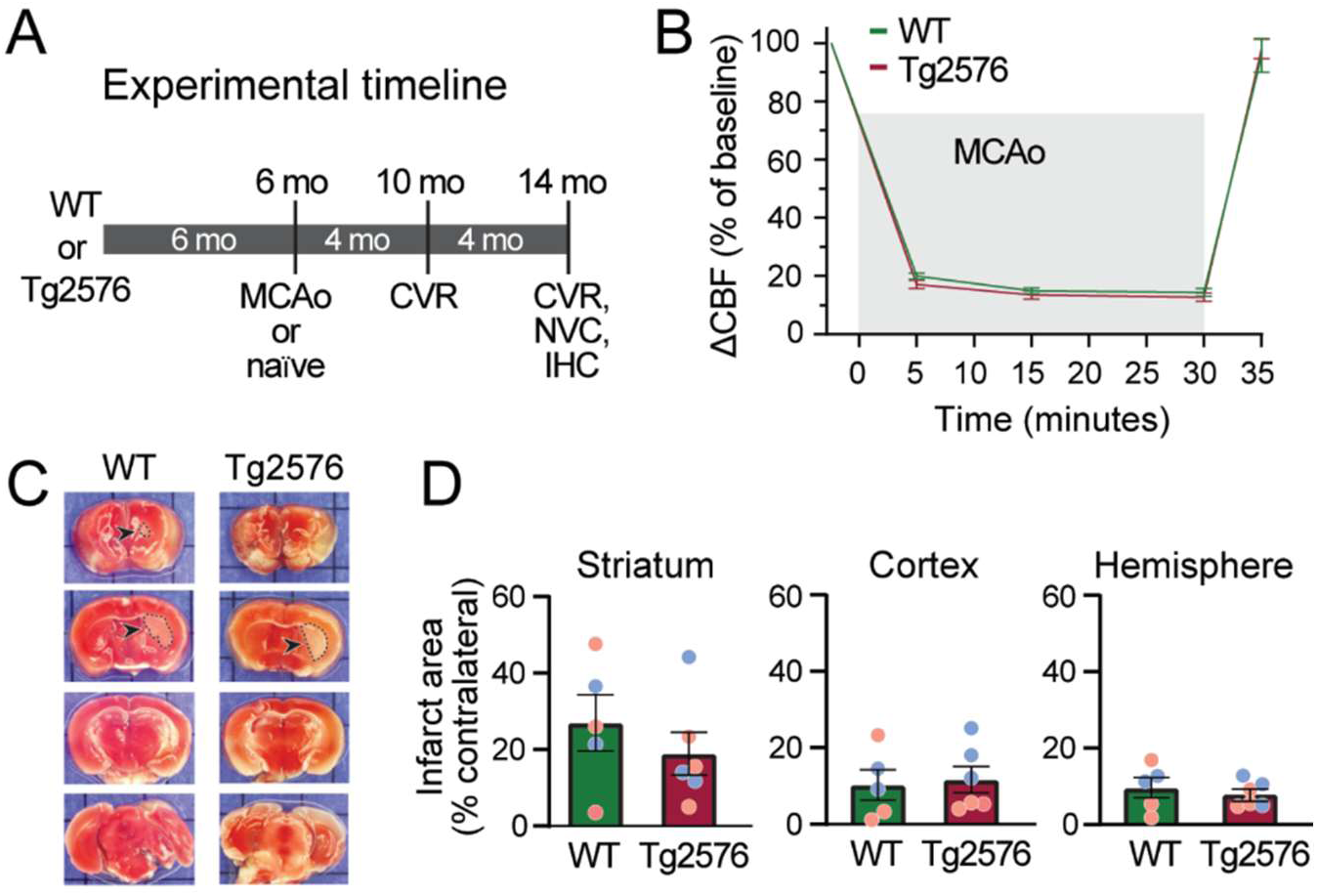
Mild subcortical ischemia induces a similar infarct in WT and Tg2576 mice. **(A)** Experimental timeline of tMSCI via middle cerebral artery occlusion (MCAo) and assessments used to evaluate disease outcomes: cerebrovascular reactivity (CVR), neurovascular coupling (NVC), and immunohistochemistry (IHC). **(B)** Change in cerebral blood flow (CBF) as a percentage of baseline during the 30-min MCAo procedure in WT and Tg2576 mice. **(C)** Example brain section stained with 2,3,5-triphenyltetrazolium chloride (TTC) 24 hours following a 30-min MCAo in WT and Tg2576 mice. Infarcted tissue does not take up TTC and remains white (dotted outline). **(D)** Quantification of infarct size displayed as a percentage of the contralateral hemisphere. Data were compared using unpaired t-tests; blue dots are from male mice (n=3 per group); orange dots are from female mice (n=3 per group).

### 2.2 Experimental timeline

Animals were assigned to naïve or mild ischemia groups, while aiming for equal distribution of sex in each group. At 6 mo of age, mice in the ischemia group were exposed to a surgically induced transient mild subcortical ischemia (tMSCI). Thus, four experimental groups were used: Wild-type (WT) naïve, WT tMSCI, Tg2576 naïve, and Tg2576 tMSCI. Mice underwent magnetic resonance imaging (MRI) scans to evaluate CVR longitudinally at 10 and 14 mo old. Next, a terminal procedure to assess NVC was performed using laser Doppler flowmetry (LDF) at 15 mo old, following which they were fixed by transcardiac perfusion and their brains harvested for immunohistochemical analysis (**Figure 1A**).

### 2.3 Transient mild subcortical ischemia

A short occlusion of the middle cerebral artery (MCA) was performed to induce tMSCI, following previously reported methods^17, 19, 20^. Reversible ischemia was induced by MCA occlusion via the intraluminal suture technique under isoflurane anesthesia (5% induction, 1% maintenance) with 30% oxygen. Body temperature was maintained at 37 ± 0.5°C throughout the procedure. A small incision was made in the skin above the right parietal skull (2 mm posterior, 6 mm lateral to bregma) and the laser Doppler probe placed in it (Moor Instruments, UK; 810 nm wavelength) to monitor MCA territory blood flow. A ventral neck incision was created to visualize the right common carotid artery (CCA) and its external (ECA) and internal (ICA) branches. An occluding filament consisting of a 6-0 monofilament nylon surgical suture (Doccol Corporation, Sharon, MA, USA) with a 0.22 mm thickness silicone rubber coated tip was inserted into the ECA, the ECA was pulled backwards caudally to align with ICA, and the filament was advanced gently into the ICA. The filament was advanced until the MCA branching point, when the LDF signal abruptly decreased, indicating low cortical blood flow in the MCA territory. The filament was left at this point to occlude the MCA for 30 minutes (mins), after which the filament was gently withdrawn to restore blood flow as measured by LDF (**Figure 1B**).

The neck and skull incision were both closed with sutures, all laser Doppler and temperature probes were subsequently removed, and mice monitored during initial reperfusion and following anesthesia cessation. After the procedure, mice were housed separately from one another and monitored for several hours post-operation to ensure recovery.

### 2.4 Infarct assessment

Infarct size was measured at 24 hours after MCAO in 2 mm thick coronal brain sections (four total) using 2,3,5-triphenyltetrazolium chloride staining and digital image analysis. Sections were incubated in 1.2% 2,3,5-triphenyltetrazolium chloride in saline for 15 mins at 37°C and then fixed in formalin for 24 hours. Slices were photographed, and infarcted (unstained white area) and uninfarcted (red stained area) regions were measured with Image J software and integrated across all four slices (**Figure 1C**). To account for the effect of edema, the infarcted area was estimated indirectly by subtracting the uninfarcted area in the ipsilateral hemisphere from the contralateral hemisphere and expressing infarct volume as a percentage of contralateral hemisphere (**Figure 1D**).

### 2.5 MRI – Data acquisition

All MRI experiments were performed at the Oregon Health & Science University Advanced Imaging Research Center. ASL-MRI was performed using an ultra-high field Bruker-Biospin 11.75 Tesla (Bruker Scientific Instruments, Billerica, MA, USA) small animal MR system equipped with a 10 cm inner-diameter gradient set and running ParaVision 5.1. Mice were anesthetized in one of two ways: (1) with inhalation of 1.5% isoflurane only or (2) with a combination anesthetic regimen using a sub-maximal dose of intraperitoneal injection of ketamine/xylazine (7.0 / 0.7 mg/kg) and inhalation of 0.75% isoflurane (**Figure 2**). Mice were secured in an animal cradle with head immobilization and supplied with 100% oxygen. Throughout the experiment, body temperature was monitored and maintained at 37 °C, and respiration was tracked using a physiological monitoring system (SA Instruments, Stony Brook, NY, USA). Radiofrequency transmission was performed using a 72 mm inner-diameter, 60 mm length Bruker volume resonator, while signal reception was achieved with an actively decoupled Bruker mouse head surface coil.

**Figure 2.**
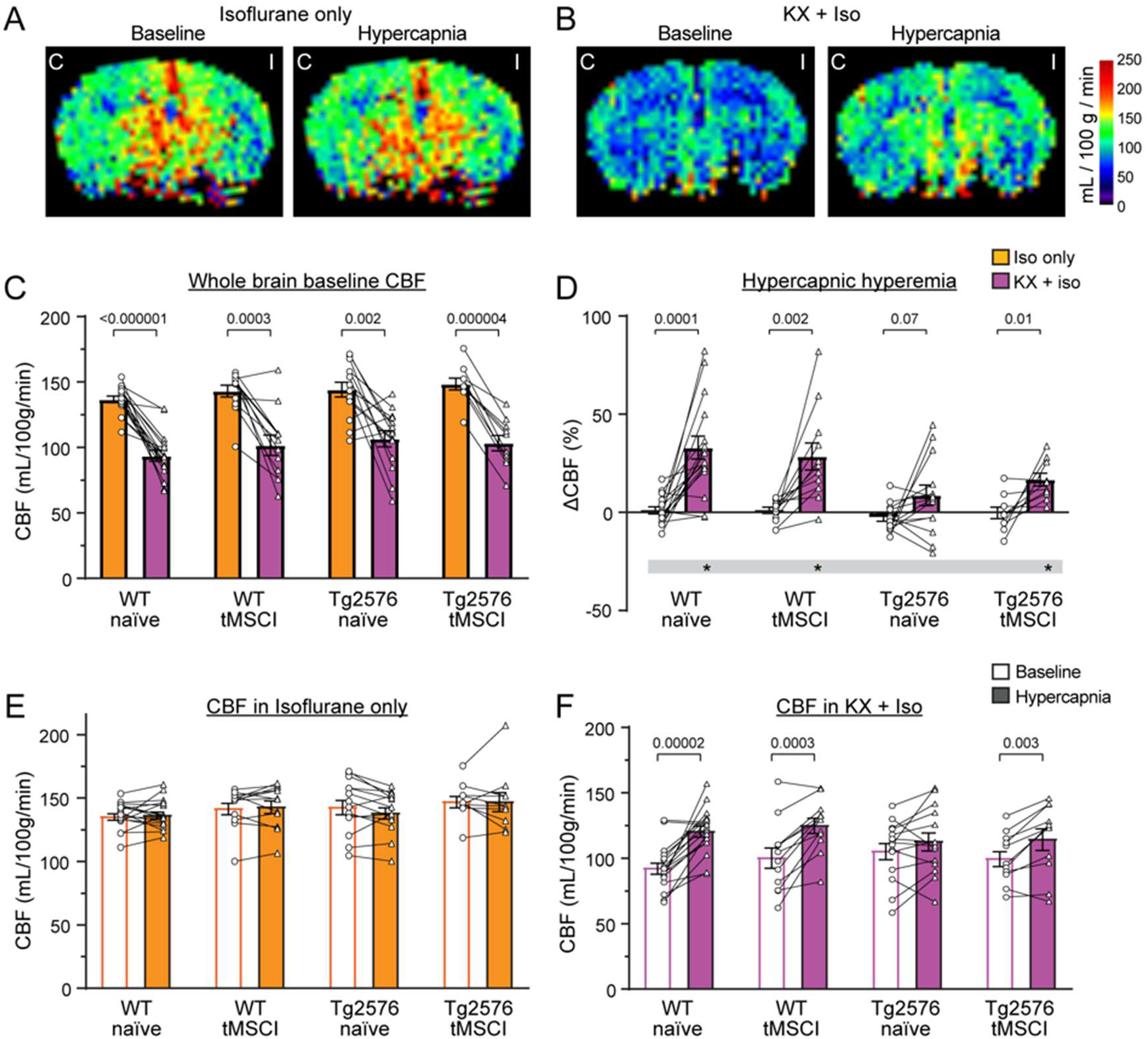
Anesthesia regimen impacts cerebrovascular flow and reactivity. **(A-B)** Representative CBF maps obtained from ASL MRI at baseline and following hypercapnia under 1.5% isoflurane **(A)** and KX + 0.75% isoflurane **(B)**; C = contralateral and I = ipsilesional. **(C)** Whole brain baseline CBF of mice in 4 experimental groups (WT naïve, WT MSCI, Tg2576 naïve, Tg2576 MSCI) under 1.5% isoflurane (circles) and KX + 0.75% isoflurane (triangles). **(D-F)** Whole brain hypercapnic hyperemia displayed as a percentage of baseline **(D)** under 1.5% isoflurane (circles) and KX + 0.75% isoflurane (triangles) and as raw CBF values **(E-F)** at baseline (circles) and following hypercapnia (triangles). Data were compared within each experimental group using paired t-tests. Asterisk depicts p<0.05 in one-sample t-tests compared to no change in CBF.

### 2.5.1 Anatomical T2-weighted acquisition

Anatomical coronal T2-weighted images were acquired approximately ∼1.6 mm posterior to the bregma using a spin-echo RARE sequence (field of view 32 mm, matrix 256 × 256, in-plane resolution 125 × 125 µm, slice thickness 0.5 mm, 25 slices, TR/effective TE = 4021/24 ms, RARE factor = 8, two signal averages).

### 2.5.2 Arterial Spin Labeling acquisition

CBF was measured using arterial spin labeling (ASL) MRI with a flow-sensitive alternating inversion recovery rapid acquisition with relaxation enhancement sequence (FAIR-RARE). Images were acquired with a recovery time of 10,000 ms, effective TE of 45 ms, echo spacing of 5 ms, and one signal average, slice thickness = 2 mm, single-slice acquisition, matrix = 128 × 128, in-plane resolution = 250 µm, RARE factor = 72, and 23 inversion times spanning 40–4400 ms. Total acquisition time for each ASL series was about 15 minutes. This approach labels inflowing arterial blood via global inversion of the equilibrium magnetization^21^. After a baseline ASL scan under 100% O2 in breathing air, mice underwent a hypercapnic challenge with exposure to 5% CO2/ 95% O2 for 10 min to allow CBF to reach a hyperemic steady state and ASL MRI performed again while maintaining hypercapnic conditions throughout the scan.

### 2.6 MRI analysis – anatomical

MRI data were processed using a custom analysis pipeline implemented with FSL (v6.0)^22^ and ANTs (v2.4.4)^23^. T2-weighted anatomical images were corrected for intensity inhomogeneity using N4 bias field correction^24^, followed by brain extraction using bet4animal.

Anatomical images were registered to a standardized mouse brain atlas^25^ using a multi-stage registration strategy consisting of rigid, affine, and nonlinear (SyN) transformations. Registration was performed using antsRegistration. Atlas label maps were subsequently transformed into individual subject space using antsApplyTransforms^26^ with nearest-neighbor interpolation to preserve discrete regional boundaries.

Anatomical volumes were calculated within each regional mask in the individual subject anatomical space. Ventricular regions were excluded in whole brain and hemispheric volume estimates.

### 2.7 MRI analysis – blood flow

Images were analyzed using the Bruker Paravision software and JIM 8.0 software (Xinapse Systems LTD, Northants, UK). Outlier value pixels (outside 2 standard deviations) were excluded, so that large arteries with high, pulsatile flow and ventricles with low flow were excluded, thus arriving at flows which consistently represent tissue microvascular flow. Ventricular regions were removed from the ASL image, guided by the structural T2 image. Mean CBF (mL/100 g/min) was quantified on this final corrected image. Major regions analyzed include the hemispheres, ACA and MCA cortical territories, hippocampus, amygdala, and thalamus on the ipsilesional and contralateral side. Selection of all regions was guided anatomically using the corresponding T2 images.

Cerebrovascular reactivity (CVR) was quantified as the percent change in CBF as a response to hypercapnia from baseline:

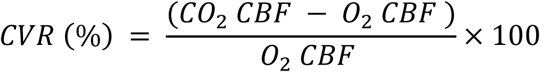

### 2.8 Laser Doppler flowmetry

Following MRI, NVC was assessed *in vivo* by quantifying hind-paw stimulation-evoked increase in blood flow in the somatosensory cortex, using LDF. Mice were anesthetized with isoflurane, and the skull was exposed, a headplate was attached, and the headplate bars were secured to a stereotaxic frame. The LDF probe was placed against the skull above the hind paw region of the somatosensory cortex (1 mm caudal of the coronal suture, 1-2 mm lateral of the sagittal suture). CBF was recorded before, during, and after mild electrical stimulation of the contralateral hind paw (2.5 mA, 3 Hz, 5 s). Three trials were performed 5 min apart, and the mean response from the trials was used for the final readout. This LDF protocol was performed on both the ipsilateral and contralateral sides of the brain with a randomly assigned order, so neither hemisphere was systematically measured first or second.

### 2.9 Laser Doppler flowmetry – analysis

LDF time-series data were analyzed using a custom Python pipeline developed for time-domain characterization of microvascular blood flow dynamics. Signals were low-pass filtered at 0.2 Hz using a fourth-order Butterworth filter applied with zero-phase forward– reverse filtering to avoid phase distortion. This filtering isolated slow hemodynamic components while suppressing high-frequency noise.

Baseline variability of the unfiltered signal was quantified as the standard deviation of flux values between 5 and 30 seconds, excluding the initial transient period. Baseline mean flow was computed from the filtered signal over the same 5–30 s window. Peak flow response was defined as the maximum filtered flux occurring between 30 and 60 s. Peak latency was calculated relative to the 30-second stimulation onset. Percent change from baseline was computed as the relative difference between baseline mean and peak value.

### 2.10 Immunohistochemistry

Immediately after LDF acquisition, a subset of mice were transcardially perfused with 1% heparinized 0.9% saline, followed by 4% paraformaldehyde (PFA). Their brains were extracted and placed in 4% PFA at 4°C overnight before being rinsed and stored in 10% PBS for subsequent analysis. Perfusion-fixed brains were paraffin-embedded and 6-μm sections were cut. Antigen retrieval was performed using standard methods (30 min incubation in citrate buffer, pH 6.0, at 80°C), and tissue sections were blocked with 3% nonfat dry milk in PBS and immunolabeled with primary antibody: rabbit anti-GFAP at 1:500 (Cat. No. 16825–1-AP, Proteintech) or 4G8 anti-amyloid antibody at 1:5000 in TBST/BSA with 0.05% azide as a preservative (Cat. No. SIG-39200, BioLegend). Appropriate secondary antibody conjugated to horseradish peroxidase was then used and reacted with 3,3’-diaminobenzidine (DAB) for visualization. Cell nuclei were lightly counterstained with hematoxylin. Whole brain sections were imaged on a Zeiss Axioscan.Z1 with a 10x/0.45NA objective and a Hitachi HV-F202 camera (resulting in a pixel size of 0.441 μm × 0.441 μm).

### 2.11 Analysis of immunohistopathology

#### 2.11.1 GFAP analysis

GFAP analysis was conducted in two cortical territories: the ACA (a region that does not experience any ischemia) and the MCA (a region that experiences transient ischemia during the tMSCI procedure). Ipsilesional hemispheres were identified by the presence of a high density of GFAP-positive astrocytes suggesting a glial scar/border in the striatal region. Regions were anatomically identified based on the Allen Brain Reference Atlas. All reasonable effort was made to match coronal sections between animals. For each region, two representative regions of interest (ROIs; 250x 250 μm^2^) were selected per animal for quantification. A cloud-based deep learning segmentation platform Biodock (AI Software Platform, Biodock 2024, Available from www.biodock.ai) was used to count the number of GFAP-positive cells. This platform was used by our group previously to train an artificial neural network (ANN) to recognize astrocytes^20^. ANN astrocyte identification was manually checked in each ROI image by a trained expert. The density of positive cells (number of cells/mm^2^) was averaged across the two ROIs for each region in each animal and reported.

#### 2.11.2 4G8 analysis

ImageJ was used to quantify 4G8-positive parenchymal Aβ plaques and depositions around parenchymal and leptomeningeal vessels. Number of plaques and vascular deposition were manually counted and classified within the cortical regions (ACA and MCA territories) of one section per animal, anatomically identified based on the Allen Brain Reference Atlas. All reasonable effort was made to match coronal sections between animals. The densities of 4G8-positive vessels and plaques were reported as the number of each normalized to the area (plaques or vessels/mm^2^). To assess stain area, H DAB color deconvolution was used to isolate the image with DAB stain, a threshold was applied (uniformly across sections), particles above 10 pixels^2^ (4.784 um^2^) were analyzed. The total stain area was normalized to the area of the cortex and reported as a percentage. The size of individual Aβ deposits was assessed using the area of each plaque detected using the same thresholding method.

### 2.12 Statistical analysis

In all figures, data are shown as mean ± SEM with the raw data values indicated by individual points and the N indicated in the bars. Paired data from the same mice at different timepoints or conditions were analyzed with paired two-sample t-tests. To analyze the effects of genotype and stroke status, two-way analysis of variance (ANOVA) with Tukey’s multiple comparisons tests were performed. The differences in data from multiple groups, regardless of genotype and stroke, were analyzed with one-way ANOVA with Tukey’s or Šídák’s multiple comparisons tests. For data calculated as an index, outcomes were analyzed for significant difference from a hypothetical value of 0 using a one-sample t-test.

## 3 RESULTS

### 3.1 Mild subcortical ischemia is induced by 30-minute MCA occlusion

We used 30-min MCA occlusion followed by reperfusion to induce transient, mild ischemia in subcortical regions in 6 mo WT and Tg2576 mice. The procedure induced a similar ∼80% decrease in blood flow in the MCA territory during occlusion (**Figure 1B**) and resulted in similar infarct sizes in the WT and Tg2576 mice (**Figure 1C-D**). We employed this tMSCI model for the entire study.

### 3.2 Anesthesia regimen impacts CBF and CVR

We performed ASL-MRI to assess CBF and CVR in 10 mo mice (∼4 mo after tMSCI) under ∼1.5% isoflurane anesthesia but found no hyperemic response to hypercapnic challenge in any group. This was accompanied by a high baseline flow measurement, prompting us to consider the potential confound of isoflurane-induced dilation masking any response. To examine this, we assessed CBF and CVR again a second time, performed one week later under a combination anesthesia using sub-maximal ketamine/xylazine (KX; 7.0 / 0.7 mg/kg) and isoflurane (0.75%; **Figure 2**), as low KX dose can induce an extended surgical plane of anesthesia at lower isoflurane concentrations^27^. When isoflurane was used alone, we found 1.5% concentration was the minimum dose necessary to prevent movement artifacts and avoid confounds in the ASL perfusion data. In the combination regimen, although the dose of each anesthetic would not be effective on its own, the additive effects of 7.0 / 0.7 mg/kg and 0.75% isoflurane were effective in sedating the mice and minimizing motion during MRI, while reducing side effects.

KX + isoflurane combination anesthesia significantly reduced baseline CBF and revealed the CVR response in all groups compared to isoflurane-only anesthesia (**Figure 2**). On average, regardless of genotype or stroke status, baseline CBF under isoflurane alone was 41.6% higher than under KX + isoflurane (p < 0.0001). CBF was significantly increased by hypercapnic challenge under KX + isoflurane combination anesthesia (WT naïve, p < 0.001; WT MSCI, p = 0.0017; Tg2576 naïve, p > 0.01; Tg2576 MSCI, p = 0.005; **Figure 2D, F**), while essentially absent under isoflurane only (all groups, p > 0.01; **Figure 2D-E**). This pattern was present in all regions quantified, including the ACA and MCA cortex territories, hippocampus, amygdala, and thalamus (**Supplemental Figure 1**). Together, these results indicate that the choice of administered anesthesia can significantly confound assessments of vascular functionality. Isoflurane, even at moderate doses, can raise baseline blood flow and mask CVR. Therefore, we use the low dose KX + low isoflurane (0.75%) combination anesthesia to assess blood flow in all subsequent experiments.

### 3.3 Volume of subcortical structures is mildly reduced after tMSCI

To examine any tissue loss induced by tMSCI, we quantified the volume of several brain regions using the T2-weighted anatomical whole brain images acquired 8 mo after tMSCI (mice aged 14 mo). We quantified the whole hemisphere, cortex, striatum, ventricles, hippocampus, amygdala, and thalamus regions in each hemisphere. Normalizing the ipsilesional volume to the contralateral volume within each animal allowed us to capture even small changes due to tissue loss (**Figure 3** and **Supplemental Figure 2**). We observed a small decrease in the volume of the striatum (**Figure 3D**) and thalamus (**Supplemental Figure 2C**) in both WT (5.03 and 5.53% reduction, respectively) and Tg2576 mice (7.59 and 2.67% reduction, respectively) exposed to tMSCI. A reduction in total cortical volume was only detected in the Tg2576 mice with tMSCI (7.56% reduction; **Figure 3B**). No differences were observed in the volume of the total hemisphere, hippocampus, amygdala, or ventricles in either genotype. These data indicate that the stroke induced by tMSCI procedure was indeed mild and limited to subcortical regions in WT mice but also spread into the cortex in the Tg2576 mice.

**Figure 3.**
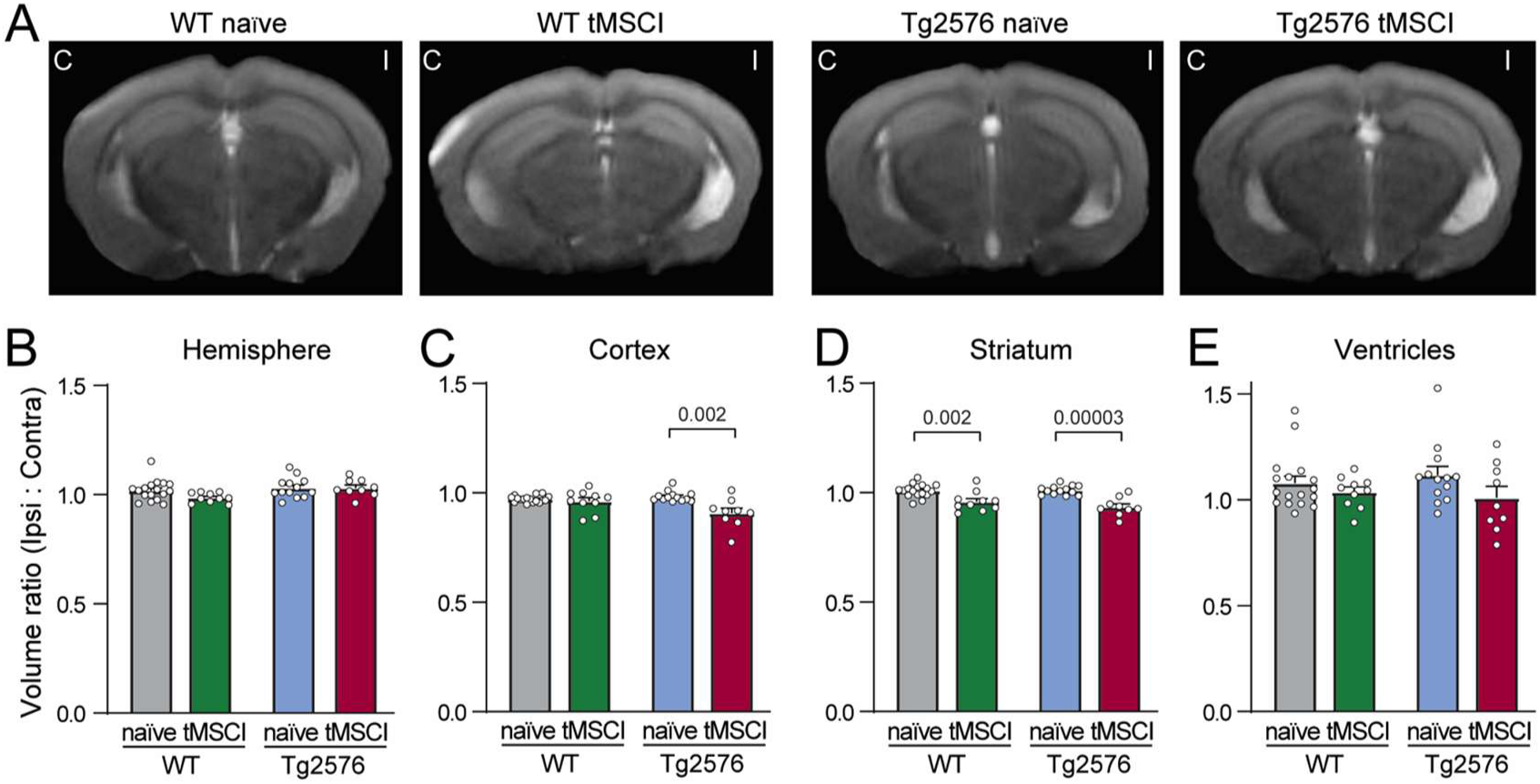
tMSCI reduces striatal volume in both genotypes but cortical volume only in Tg2576 mice. **(A)** Representative MRI T2-weighted images for each experimental group; C = contralateral and I = ipsilesional. **(B-E)** Volume of anatomical regions displayed as a ratio of ipsilesional to contralateral for hemisphere **(B)**, cortex **(C)**, striatum **(D)**, and ventricle **(E)**. Data were analyzed with one-way ANOVAs followed by comparison of naïve and tMSCI groups within each genotype with Šídák’s multiple comparisons test.

### 3.4 Cerebrovascular reactivity is differentially altered by age and insult

We quantified CBF using ASL-MRI 4 mo and 8 mo after tMSCI in the following regions: the whole hemispheres, the ACA and MCA territories of the cortex, hippocampus, amygdala, and thalamus. We found no differences between groups in the resting level of CBF at either timepoint (**Supplemental Table 1**). When compared to baseline CBF, hypercapnia-induced CBF was significantly higher in the WT naïve and WT tMSCI mice in all regions quantified (**Figure 4** and **Supplemental Figures 3-4**). The Tg2576 naïve mice showed a lack of CVR in all regions except a small increase in the ipsilesional hippocampus (**Figure 4** and **Supplemental Figure 3A and C**). Surprisingly, the Tg2576 mice exposed to tMSCI showed a small but significant CVR response in all regions except the contralateral amygdala, which was trending but not significant (**Figure 4** and **Supplemental Figure 3B**). When the CVR index (hypercapnia-induced change in CBF) was compared between groups, we observed minimal effects of tMSCI on WT mice at the 4 mo post-tMSCI timepoint. Tg2576 mice exposed to tMSCI tended to show consistently smaller responses compared to WTs regardless of stroke status at this timepoint (**Supplemental Figure 4**). Although CVR generally tended to be lower in the Tg2576 naive group than WT naïve controls, they showed a statistical decrease only in the ipsilesional thalamus (**Supplemental Figure 4F**). At 4 months post-tMSCI, one-sample t-tests indicated that there was a significant CVR response above baseline flow in WT naïve, WT tMSCI and Tg2576 tMSCI groups in all regions except the contralateral (left) amygdala and MCA territory. In contrast, in the Tg2576 naïve mice, CVR was significant only in the thalamus and contralateral hippocampus.

**Figure 4.**
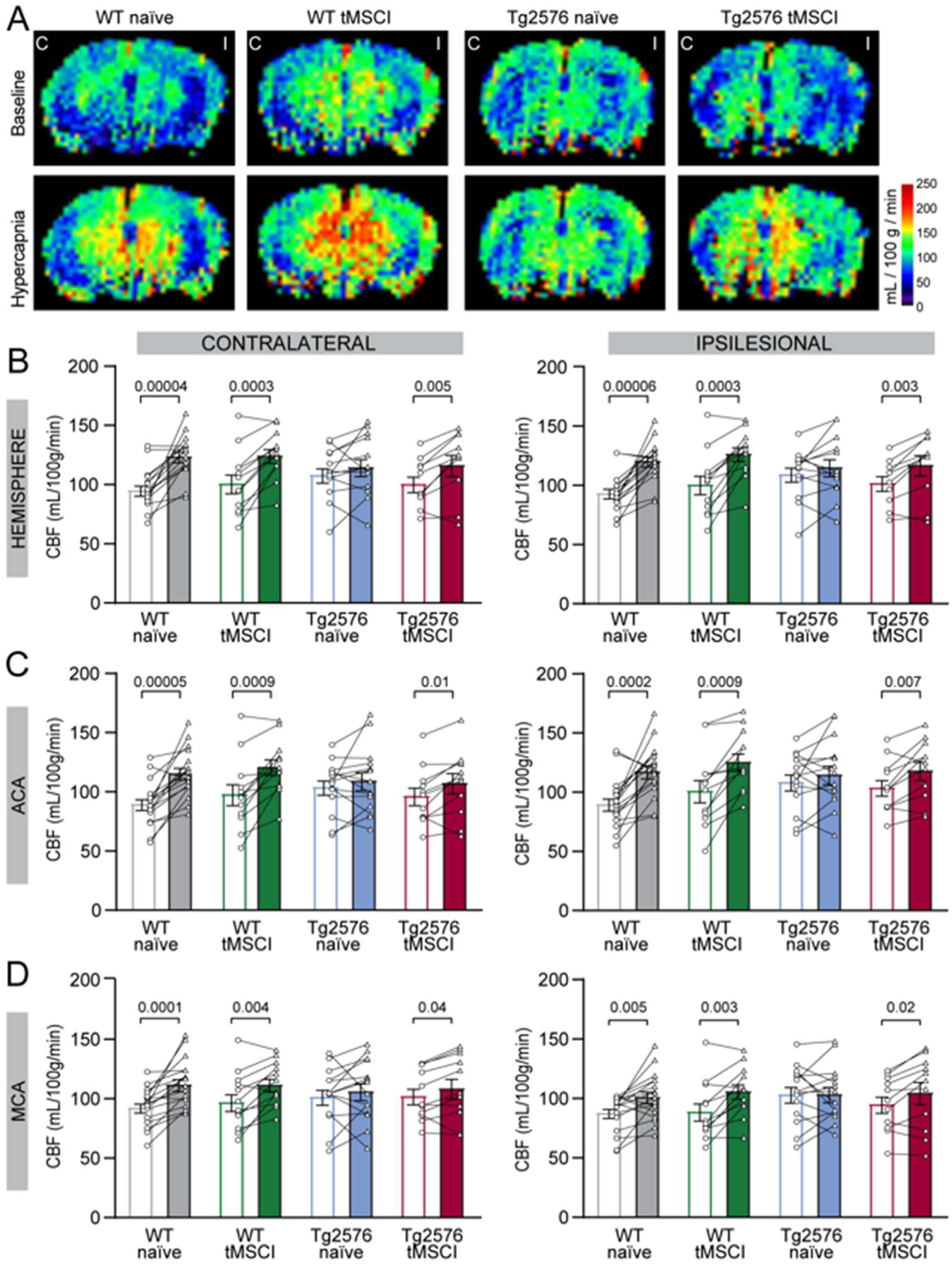
Cerebrovascular reactivity is impaired by Aβ but not stroke at 4 mo post-tMSCI. **(A)** Representative CBF maps obtained from ASL MRI at baseline (above) and following hypercapnia (below) at 4 mo post-tMSCI at 10 mo old; C = contralateral and I = ipsilesional. **(B-D)** Hypercapnic hyperemia displayed as raw CBF values at baseline (circles) and following hypercapnia (triangles) for contralateral and ipsilateral hemisphere **(B)**, ACA **(C)**, and MCA **(D)** territories. Data were compared within each experimental group using paired t-tests.

At the 8 mo post-tMSCI timepoint (14 mo of age; **Figure 5** and **Supplemental Figure 5, 6**), WT tMSCI and Tg2576 naïve groups showed loss of hypercapnia-induced hyperemia bilaterally in both ACA and MCA regions (**Figure 5C, D**) as well as the ipsilesional hippocampus (**Supplemental Figure 5A**), but not the Tg2576 tMSCI group. The amygdala displayed a bilateral lack of hyperemia in both WT tMSCI and Tg2576 tMSCI groups (**Supplemental Figure 5B**), while the thalamus showed a robust response in all groups (**Supplemental Figure 5C**). At this timepoint (8 mo post-tMSCI), the CVR index (**Supplemental Figure 6**) was statistically significant (above baseline) in all regions only in WT naïve mice, while a lack of response was observed in most regions of the WT tMSCI and Tg2576 naïve mice. The Tg2576 tMSCI mice showed mixed results, with the CVR being robust in the hemispheres, hippocampus, ACA cortex and thalamus, but weaker in the amygdala and MCA cortex (**Supplemental Figure 6**). There were no statistical differences between groups in the CVR index aside from a decrease in the thalamus of Tg2576 naïve group compared to the WT naïve group (**Supplemental Figure 6F**).

**Figure 5.**
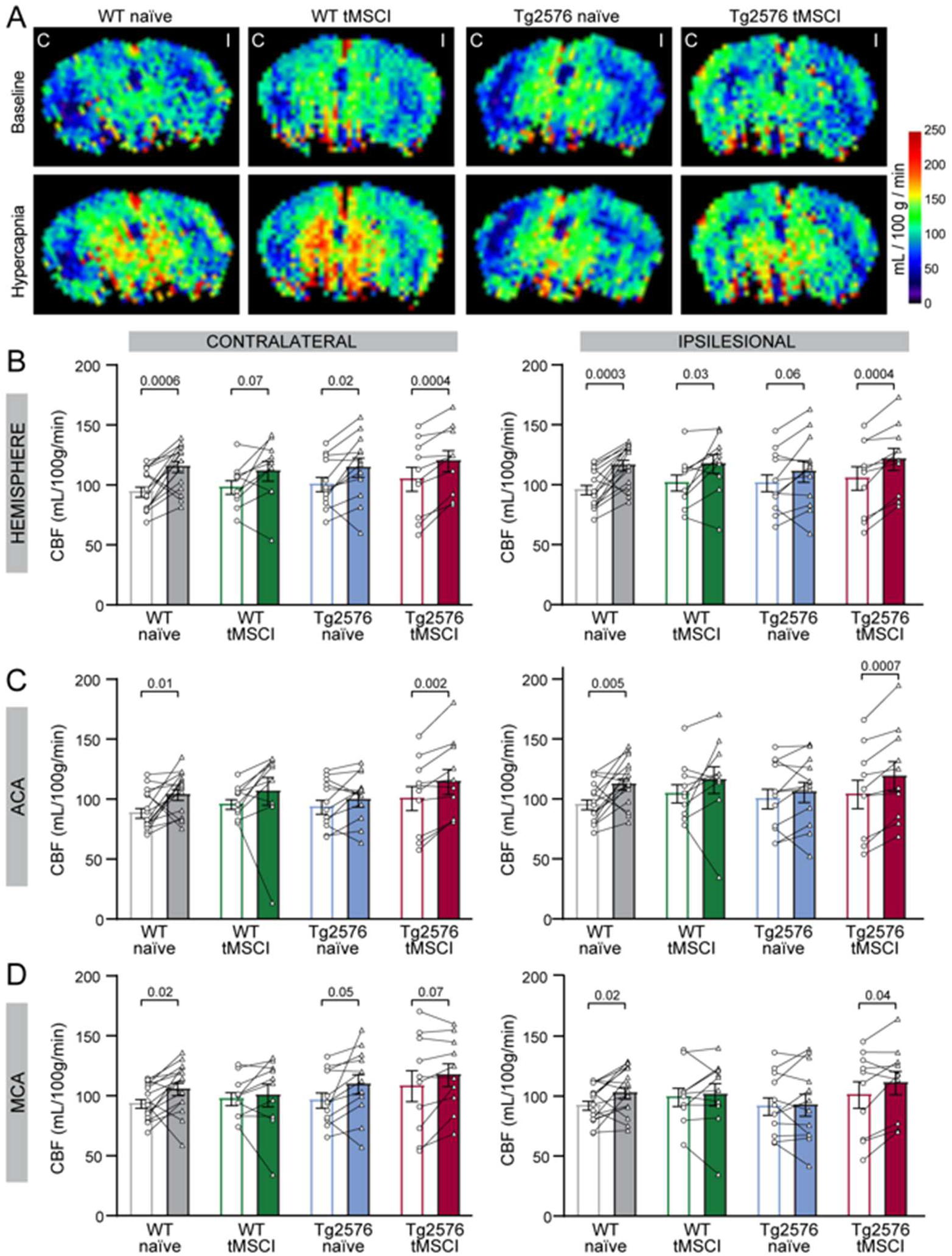
Cerebrovascular reactivity is impaired by Aβ or stroke separately at 8 mo post-tMSCI but retained when Aβ and stroke coexist. **(A)** Representative CBF maps obtained from ASL MRI at baseline (above) and following hypercapnia (below) at 8 mo post-tMSCI at 14 mo old; C = contralateral and I = ipsilesional. **(B-D)** Hypercapnic hyperemia displayed as raw CBF values at baseline (circles) and following hypercapnia (triangles) for contralateral and ipsilateral hemisphere **(B)**, ACA **(C)**, and MCA **(D)** territories. Data were compared within each experimental group using paired t-tests.

As the 4 mo and 8 mo post-tMSCI measurements were performed in the same mice longitudinally (except a few that did not survive to the second timepoint), we further examined whether any consistent change in the CVR response occurs over this timeframe of aging. We measured a worsening of the CVR index in the ipsilesional hemisphere and the MCA territory of WT tMSCI mice from 4 mo to 8 mo timepoint (**Figure 6A, C**). These data are in accordance with our ischemia model, where the MCA region transiently lacks blood flow during occlusion. WT tMSCI mice also showed a trending decrease in CVR in the ACA (**Figure 6B**) and amygdala (**Supplemental Figure 7B**), but no statistical changes were observed in other regions over time.

**Figure 6.**
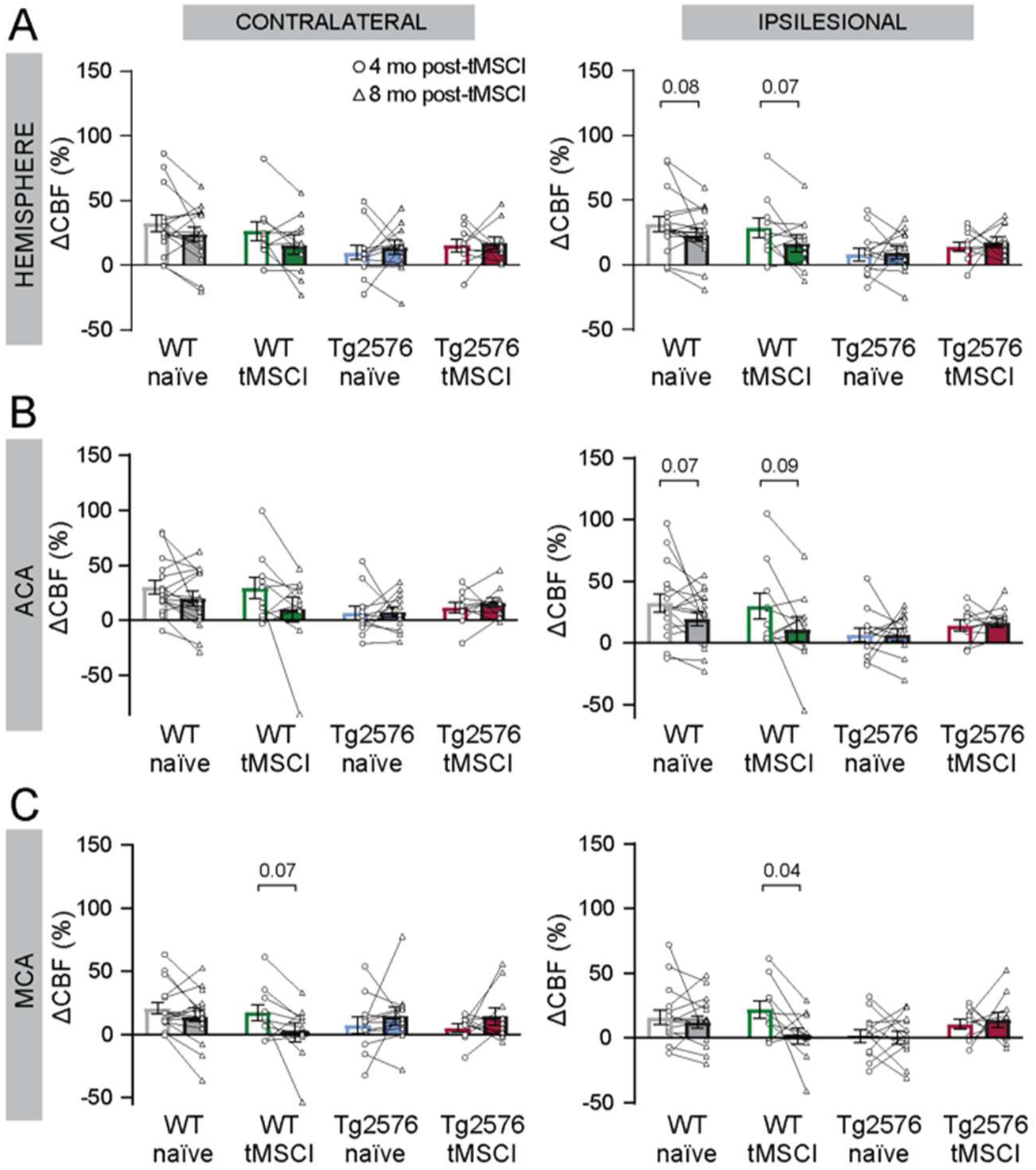
Longitudinal analysis shows that cerebrovascular reactivity diminishes over time in the MCA region after subcortical stroke from 4 to 8 mo post-tMSCI. **(A-C)** Hypercapnic hyperemia displayed as a percentage of baseline at 4 mo post-tMSCI (circles) and 8 mo post-tMSCI (triangles) for contralateral and ipsilateral hemisphere **(A)**, ACA **(B)**, and MCA **(C)** territories. Data were compared over time within each experimental group using paired t-tests.

### 3.5 Neurovascular coupling is impaired by ischemia

Following MRI (at 9 mo post-tMSCI), we subjected the same groups of mice to LDF under anesthesia to measure changes in somatosensory CBF in response to hindpaw stimulation as a measure of NVC (**Figure 7**). We observed an impairment of NVC only in the WT tMSCI group, in which both the ipsilesional response magnitude was lower and the time to reach peak was longer. WT tMSCI mice showed a similar trend in the contralateral hemisphere, but it did not reach significance. Surprisingly, at this age, neither the Tg2576 naïve nor the Tg2576 tMSCI mice showed an impairment in NVC compared to WT naïve controls. However, the magnitude of NVC was statistically lower in the ipsilesional hemisphere in Tg2576 tMSCI mice when compared to the contralateral hemisphere.

**Figure 7.**
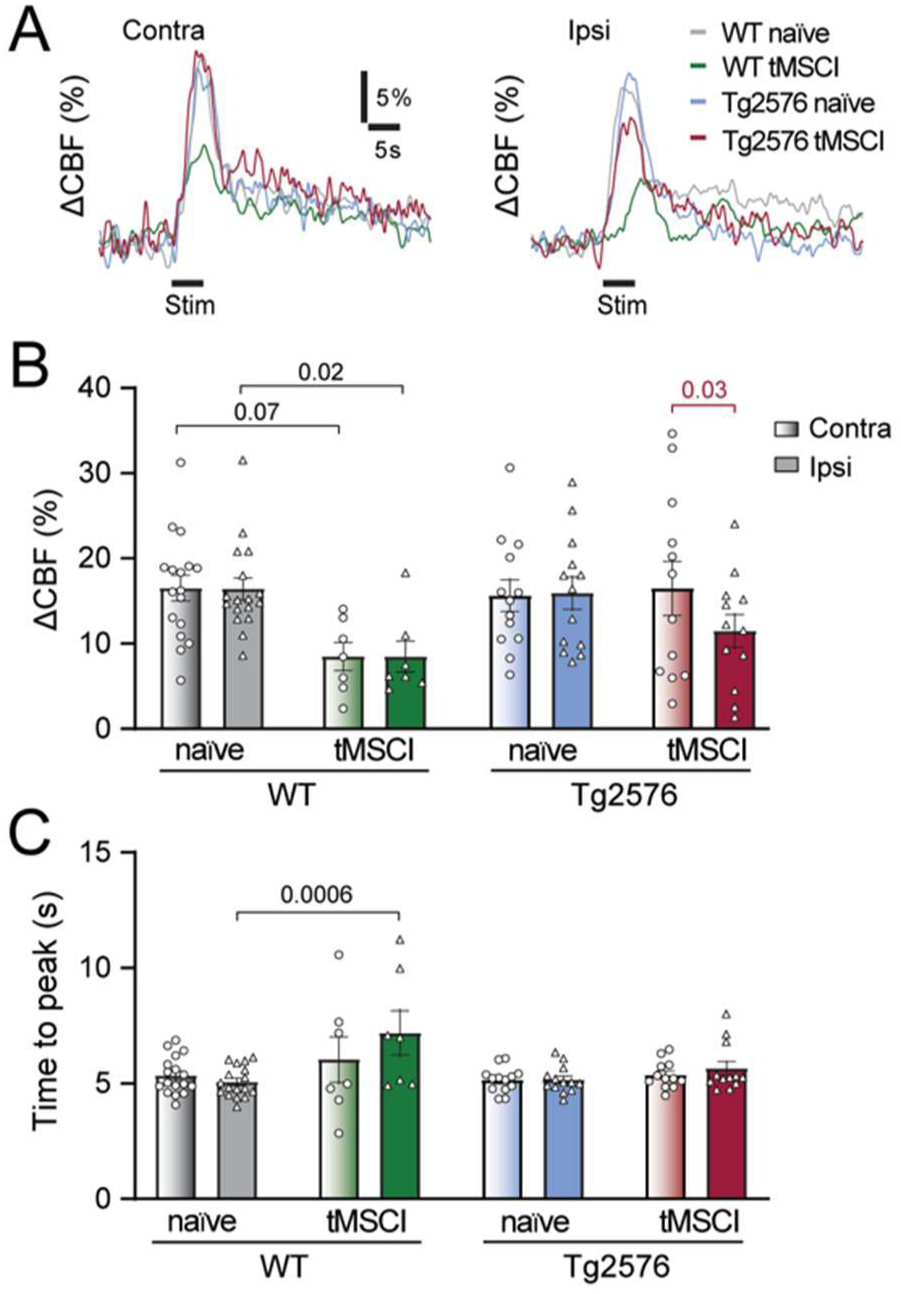
Neurovascular coupling is impaired bilaterally by subcortical stroke at 9 mo post-tMSCI, but only ipsilesionally when stroke occurs in presence of Aβ. **(A)** Representative laser Doppler flowmetry traces of blow flow response to hindpaw stimulation in the contralateral and ipsilesional somatosensory cortex (indicated by gradient and solid bars, respectively). **(B)** Peak blood flow response to stimulation quantified as a percentage of baseline. **(C)** Time to reach peak response from onset of stimulation. Data were analyzed with one-way ANOVAs, followed by comparison of naïve and tMSCI groups within each genotype with Šídák’s multiple comparisons test (black), and with unpaired t-tests within each group comparing contralateral to ipsilesional values (red).

### 3.6 Ischemia induces persistent reactive astrocytes and enhances Aβ deposition

As astrocytes modulate NVC under physiological conditions^28^ and have been shown to impair it when they become reactive, especially after stroke^17^, we next sought to quantify the level of reactive astrogliosis in each group. We quantified the number of GFAP+ astrocytes in the ACA and MCA regions (**Figure 8A-D**). As expected, the number of GFAP+ astrocytes was higher in Tg2576 mice in all regions compared to WTs, except in the ipsilesional MCA territory of WT tMSCI mice (**Figure 8B, D**). In this region, the higher number of GFAP+ astrocytes was comparable to that observed in Tg2576 mice (**Figure 8D**).

**Figure 8.**
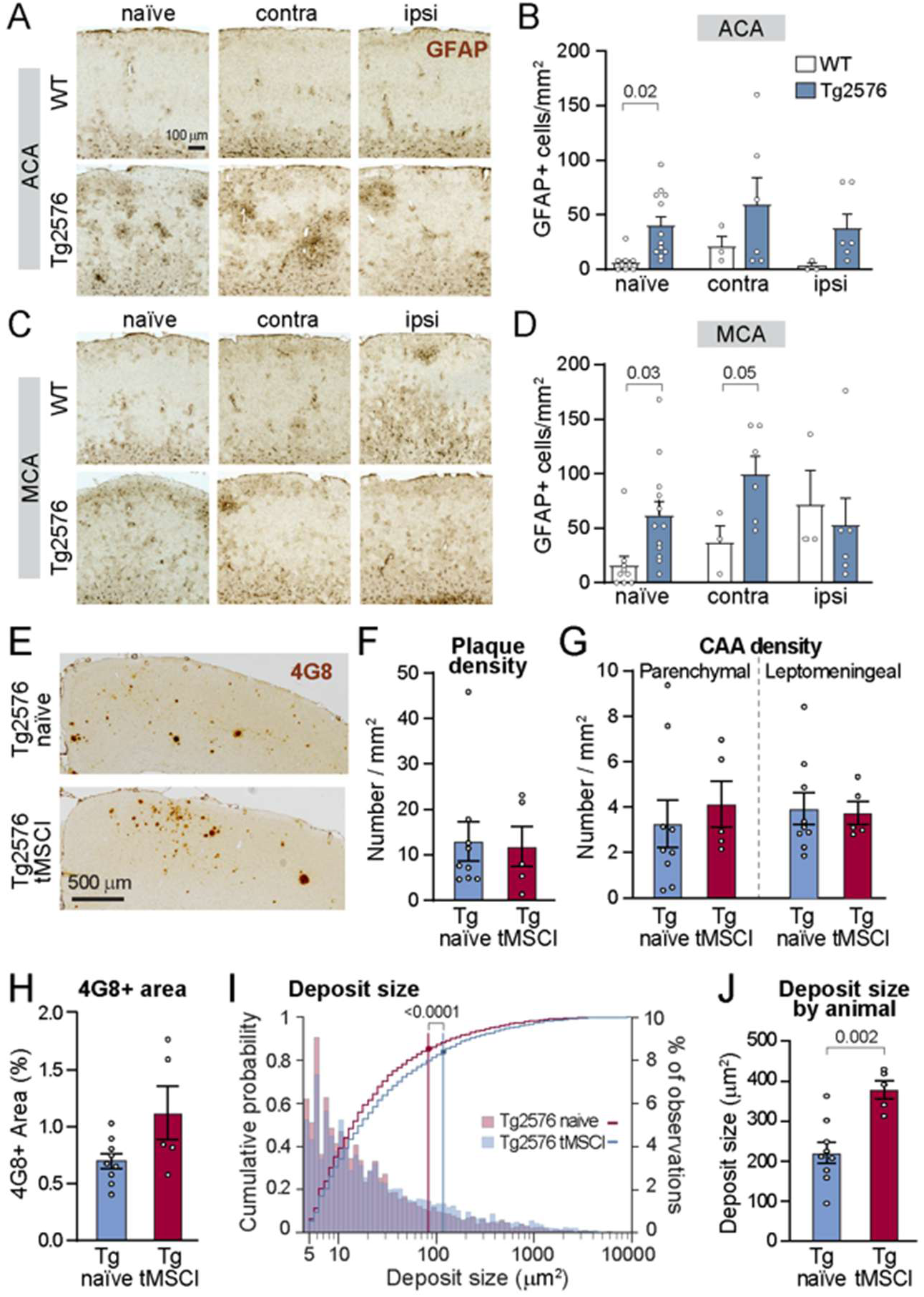
Reactive astrocytes persist and Aβ deposition is enhanced 9 mo post-tMSCI. **(A & C)** Representative GFAP labeling in the anterior cerebral artery (ACA) territory **(A)** and MCA territory **(C)** of cortical sections from WT and Tg2576 naïve and tMSCI mice. **(B & D)** Number of cells positive for GFAP in WT and Tg2576 mice in naïve, contralateral, and ipsilesional ACA **(B)** and MCA **(D)** regions. (WT naïve n=8; Tg2576 naïve n=12; WT tMSCI n=3; Tg2576 tMSCI n=6) **(E)** Representative images of cortical sections stained with 4G8 from Tg2576 naïve (top; n=9) and Tg2576 tMSCI (bottom; n=5) mice. **(F-G)** Number of 4G8-positive deposits quantified as the density of neuropil plaques **(F)** and cerebral amyloid angiopathy (CAA) of parenchymal and leptomeningeal origin **(G)**. **(H)** Percent of the cortical area positive for 4G8. **(I)** The size of 4G8-positive Aβ deposits displayed as a logarithmic histogram overlaid with cumulative probability. **(J)** The size of 4G8-positive deposits pooled by mouse **(J)**. Data were analyzed with unpaired t-tests for normally distributed data or Mann Whitney tests if a Shapiro-Wilk test normality test revealed the data was non-normal.

Given the literature that ischemia induces altered Aβ processing in rodents^18, 29–31^, albeit transiently and largely only demonstrated in rats, we wondered whether tMSCI would increase Aβ burden in the Tg2576 mouse model. Thus, we used 4G8 immunolabeling to compare Aβ deposition in Tg2576 naïve and tMSCI groups (**Figure 8E-J**). We found no differences between the number of plaques deposited in the neuropil (**Figure 8F**), nor in the density of cerebral amyloid angiopathy in parenchymal or leptomeningeal vessels (**Figure 8G**). However, the percentage of tissue area positive for 4G8 appeared to be higher (**Figure 8H**) and the size of the Aβ deposits was significantly larger (**Figure 8I-J**) in the Tg2576 mice exposed to tMSCI compared to naïve Tg2576 mice. This indicates that there is indeed an increase in Aβ accumulation and deposition in Tg2576 mice after a mild stroke.

## 4 DISCUSSION

In this study, we sought to investigate how mild ischemia contributes to ADRD pathology through cerebrovascular dysfunction. To accomplish this, we developed an innovative mixed pathology model using Tg2576 AD-like mice and a mild subcortical stroke. In this model, we characterized changes in vascular function and histopathology at chronic time points after the ischemic insult. First, we established an anesthesia regimen to accurately assess vascular function. We discovered that impairment of CVR is impacted by Aβ accumulation more strongly at earlier ages and ischemia at later ages. We also found that NVC dysfunction is associated with ischemia more than Aβ. Surprisingly, Tg2576 mice exposed to ischemia did not show significant impairments in either CVR or NVC outcomes, compared to the impairments observed in WT mice with stroke or naïve Tg2576 mice at the middle-age timepoints we examined. Mild ischemia increased reactive astrocytes up to 8 mo later in WT mice. Tg2576 naïve mice had higher numbers of GFAP immunoreactive astrocytes and this was not further exacerbated by tMSCI. Despite the lower burden of cerebrovascular impairments, we observed larger Aβ deposit size in Tg2576 mice exposed to mild ischemia.

In ASL MRI experiments, the main purpose of anesthesia is to minimize movement of the animals to produce a clear image without motion artifacts. Isoflurane is a commonly used inhalant anesthetic that allows for more precise control over the depth of anesthesia and consciousness but causes vasodilation^32^, suppresses neuronal activation^33^, and astrocyte calcium signaling^34^. KX is an anesthetic duo that reduces heart rate, causes analgesia, and induces paralysis^35^. In mice, ketamine and xylazine also have short half-lives of approximately 13 min and 1-2 h, respectively^27^. KX can prolong the surgical plane of anesthesia at lower isoflurane concentrations, without the need for supplementary injections of KX or increased isoflurane, even beyond the timepoint at which KX has likely been metabolized^27^. Therefore, combination KX + 0.75% isoflurane is an effective method of anesthesia to administer to mice undergoing MRI scans, especially if the goal is to capture perfusion measurements.

It was interesting that the reduction in CVR was evident in the Tg2576 naïve group at 10 mo of age, whereas it was more pronounced in the WT tMSCI group at 14 mo of age (8 mo after tMSCI). Yet, at both timepoints, it appears that the confluence of mild stroke and Aβ in the Tg2576 tMSCI group led to normal CVR responses in essentially all regions at both timepoints, except the amygdala at the later age. Future studies should confirm this observation and examine the mechanisms underlying this potential interaction.

Our results also highlight heterogeneity of CVR response and its vulnerability in different brain regions (**Supplemental Figure 4 and 6**), despite a similar baseline CBF across regions (**Figure 3 and 4**). CVR in the cortex was generally lower even in WT controls compared to other brain regions, and it was most severely impaired in Tg2576 mice.

Additionally, while the hippocampus, amygdala and thalamus all have similar baseline CBF, the hippocampus has large CVR responses yet is sensitive to disruption; the amygdala has lower CVR responses and also is sensitive to disruption; and the thalamus has large CVR responses and, although reduced in size, it is more resistant in the face of stroke or Aβ (change in CBF significantly above baseline in all conditions). Intriguingly, this vulnerability of CVR reflects the progression of Thal staging in human patients with AD, where cortical Aβ pathology is observed first (phase 1), followed by the hippocampus and amygdala (phase 2), and occurs later in the thalamus (phase 3)^36, 37^. This is especially noteworthy as Aβ pathology progression in the Tg2576 mouse model closely follows human Thal staging^38^. The concordance of regional Aβ spread and CVR impairment could indicate the early direct impact of Aβ on microvascular tone, as previously reported^39^. Whether this spatial overlap in the progression Aβ pathology and vascular dysfunction induced by subcortical stroke synergize to impair cognitive behavior outcomes remains to be determined.

Our study revealed that even a small stroke impairs NVC responses, with the magnitude being lower and the delay-to-peak being longer in the WT tMSCI group. Although the combination of mild stroke and Aβ (Tg2576 tMSCI group) led to lower NVC in the ipsilesional hemisphere compared to within animal contralateral hemisphere, it surprisingly was not statistically different from naïve mice of either genotype. This indicates that NVC dysfunction is mild and restricted to the stroke hemisphere in Tg2576 mice, at least at this middle-age timepoint (15 mo age). The differences in CVR and NVC observed in our study indicate divergent underlying mechanisms. CVR depends mainly on endothelial cells^40–42^ with some contribution from astrocytes^43^, while NVC depends on signals from neurons and astrocytes^44, 45^. The stronger impact of stroke on NVC suggests potential astrocyte signaling dysfunction after ischemia, while the earlier loss of CVR in Tg2576 mice suggests an endothelial or vascular-intrinsic dysfunction due to Aβ. Such cell-specific mechanisms and how they interact will need to be the focus of future studies.

GFAP staining showed increased reactive astrocytes in Tg2576 naïve mice compared to WT naïve mice, as previously demonstrated^46^. We show here that, even 8 mo after a mild stroke, reactive astrocytes were increased in the ipsilesional MCA territory of WT mice and matched that observed in the Tg2576 mice. We also found that the average Aβ deposit size was larger in Tg2576 mice exposed to mild ischemia compared to naïve Tg2576s, although no differences were observed in the density of cortical Aβ plaques in the neuropil or perivascular deposits. Larger plaque sizes can indicate a merging and clustering of plaques over time as disease progresses^47^. Thus, our results indicate that ischemia accelerates Aβ deposition.

In conclusion, we have established and characterized a novel mixed model of AD-like disease in which a mild subcortical ischemia precedes deposition of Aβ. Notably, we have shown that long term pathological effects of stroke persist even 8-9 mo after ischemia and that there is considerable variability in the timeline and regional progression of the different types of vascular impairments (CVR vs NVC) driven by ischemia and Aβ. Despite vascular function being impaired in the groups with either mild stroke or Aβ alone (WT tMSCI and Tg2576 naïve, respectively), we surprisingly observed that mice with mild stroke and Aβ (Tg2576 tMCSI) counterintuitively had better vascular function. Indeed, our results suggest that Aβ may alleviate the impact of stroke on vascular function at these middle age timepoints. Nonetheless, we discovered that ischemia can enhance Aβ deposition, which may potentially accelerate disease progression. Future studies are needed to delve deeper into the temporal and spatial role of a mild subcortical ischemia in the onset and progression of AD pathology, especially at older ages. Additionally, questions arise regarding the effect that ischemia-induced cerebrovascular dysfunction has on AD-related cognitive and behavioral deficits. Further characterization of this novel model may reveal disease outcomes and the intersecting pathophysiology of early strokes and Aβ accumulation in the brain, which in turn may shed new light on molecular mechanisms and enable new therapeutic targets to abate cognitive decline caused by stroke and Alzheimer’s disease.

## ACKNOWLEDGEMENTS

We thank Steve J. Sullivan, Heather L. McConnell, Laura M. Knittel, Nicole Libal, and Victoria Krajbich for their integral contributions to this project. We also acknowledge the veterinarians & staff in the OHSU Department of Comparative Medicine for helping us care for our animals, and technical assistance by staff in the OHSU Advanced Light Microscopy shared resource (RRID: SCR_009961). This work was supported by NIH S10OD030459 for the 11.75T Bruker MRI Instrument housed in OHSU’s Advanced Imaging Research Center (RRID: SCR_009960). The research reported in this publication used computational infrastructure supported by the Office of Research Infrastructure Programs, Office of the Director, of the National Institutes of Health under Award Number S10OD034224 (RRID:SCR_009959). The content is solely the responsibility of the authors and does not necessarily represent the official views of the National Institutes of Health.

## ANIMAL STUDY APPROVAL

This study involved the utilization of animal models. All experiments were approved by the Oregon Health & Science University Institutional Animal Care and Use Committee. This study did not involve the use of non-human primates or human subjects.

## AUTHOR CONTRIBUTIONS

OI: project management, experimental design, data collection, data analysis

SSW: data collection, data analysis, figure creation, manuscript writing

BZ: data analysis

WZ: stroke surgeries, data collection, data analysis TLS: data collection

MMP: experimental design, data collection RLW: experimental design

AM: project management, experimental design, figure creation, manuscript writing

