## Supplemental Files for "Stroke and Alzheimer’s disease have distinct consequences on neurovascular function but synergize to increase amyloid deposition"

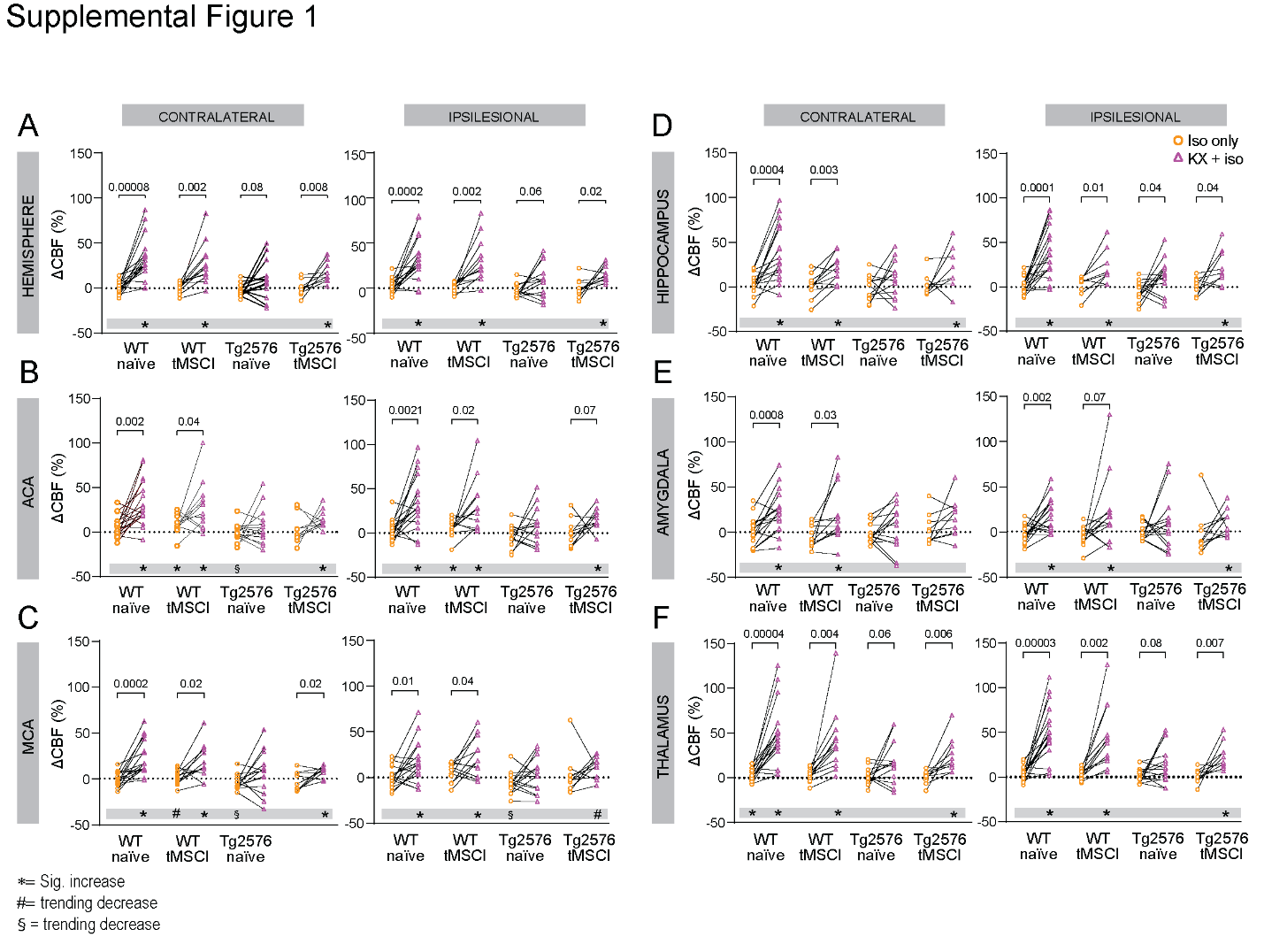

### Supplemental Figure 1. Anesthetic regimen affects blood flow across brain regions.

**(A-F)** Hypercapnic hyperemia displayed as a percentage of baseline under 1.5% isoflurane (orange circles) and KX + 0.75% isoflurane (magenta triangles) for contralateral and ipsilateral hemisphere **(A)**, ACA **(B)**, MCA **(C)**, hippocampus **(D)**, amygdala **(E)**, and thalamus **(F)**. Data were compared within each experimental group using paired t-tests. Symbols in gray bars depict results of one-sample t-tests comparing data to baseline CBF. * indicates significant increase (p<0.05), # indicates a trending increase (p<0.1), § indicates a trending decrease.

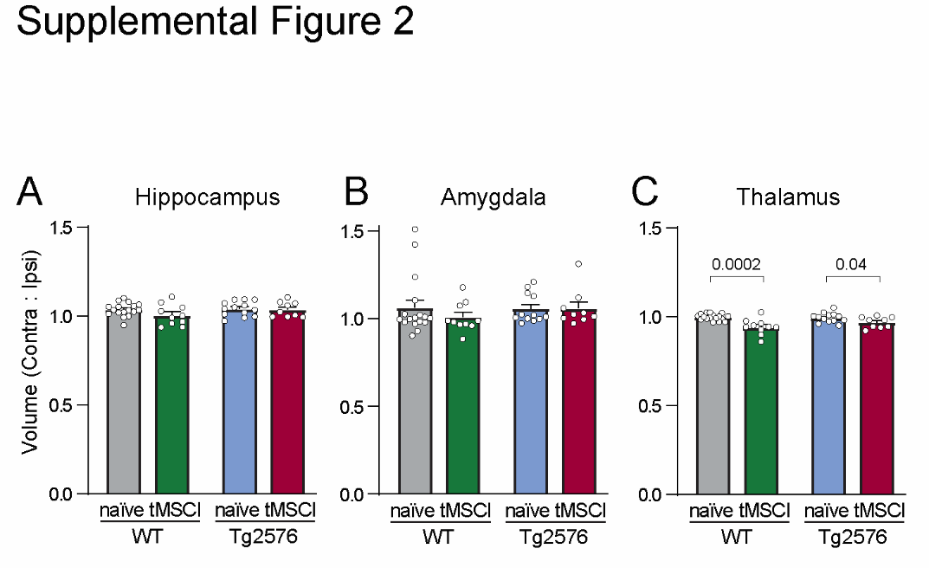

### Supplemental Figure 2. Ischemia reduces thalamic volume but not that of the hippocampus or amygdala.

**(A-C)** Volume of anatomical regions displayed as a ratio of ipsilesional to contralateral for hippocampus **(A)**, amygdala **(B)**, and thalamus **(C)**. Data were analyzed with one-way ANOVAs followed by comparison of naïve and tMSCI groups within each genotype with Šídák's multiple comparisons test.

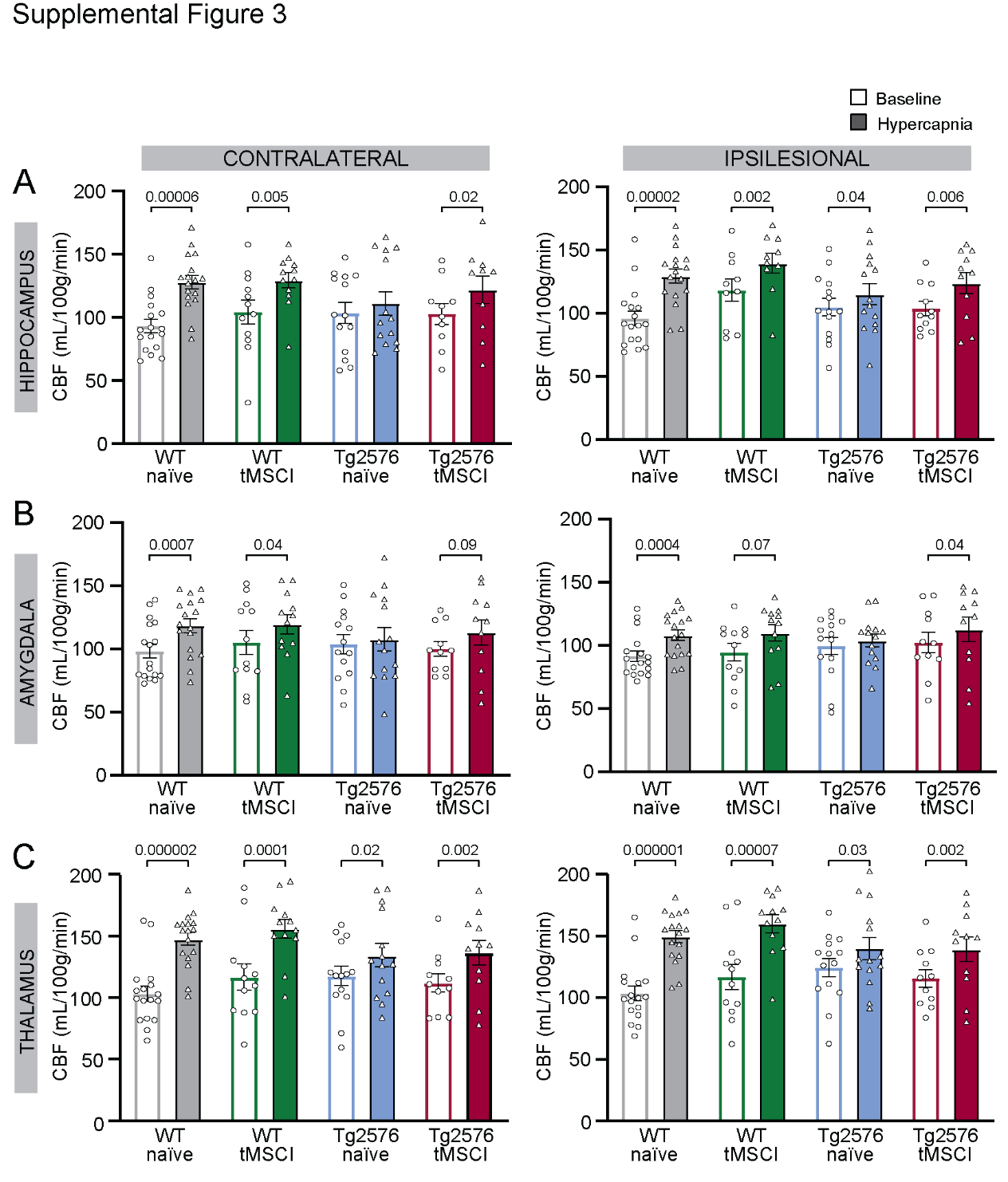

### Supplemental Figure 3. Cerebrovascular reactivity is reduced by Aβ in the hippocampus and amygdala, but not by subcortical stroke at 4 mo post-tMSCI.

**(A-C)** Hypercapnic hyperemia displayed as raw CBF values at baseline (circles) and following hypercapnia (triangles) for contralateral and ipsilateral hippocampus **(A)**, amygdala **(B)**, and thalamus **(C)**. Data were compared within each experimental group using paired t-tests.

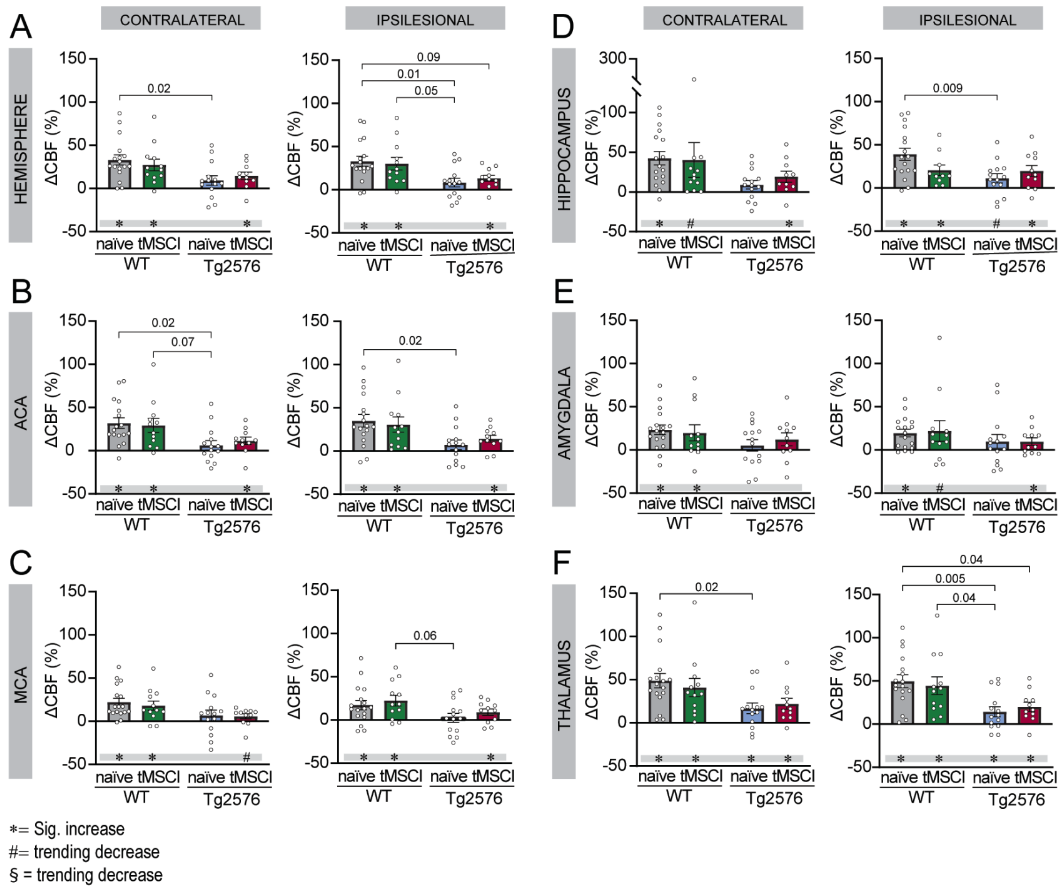

### Supplemental Figure 4. Aβ and stroke differentially affect cerebrovascular reactivity by region at 4 mo post-tMSCI.

**(A-F)** Hypercapnic hyperemia displayed as a percentage of baseline CBF for contralateral and ipsilateral hemisphere **(A)**, ACA **(B)**, MCA **(C)**, hippocampus **(D)**, amygdala **(E)**, and thalamus **(F)**. Data were compared within each experimental group using one-way ANOVAs with Tukey’s multiple comparisons. Symbols in gray bars depict results of one-sample t-tests comparing data to baseline CBF. * indicates significant increase (p<0.05), # indicates a trending increase (p<0.1).

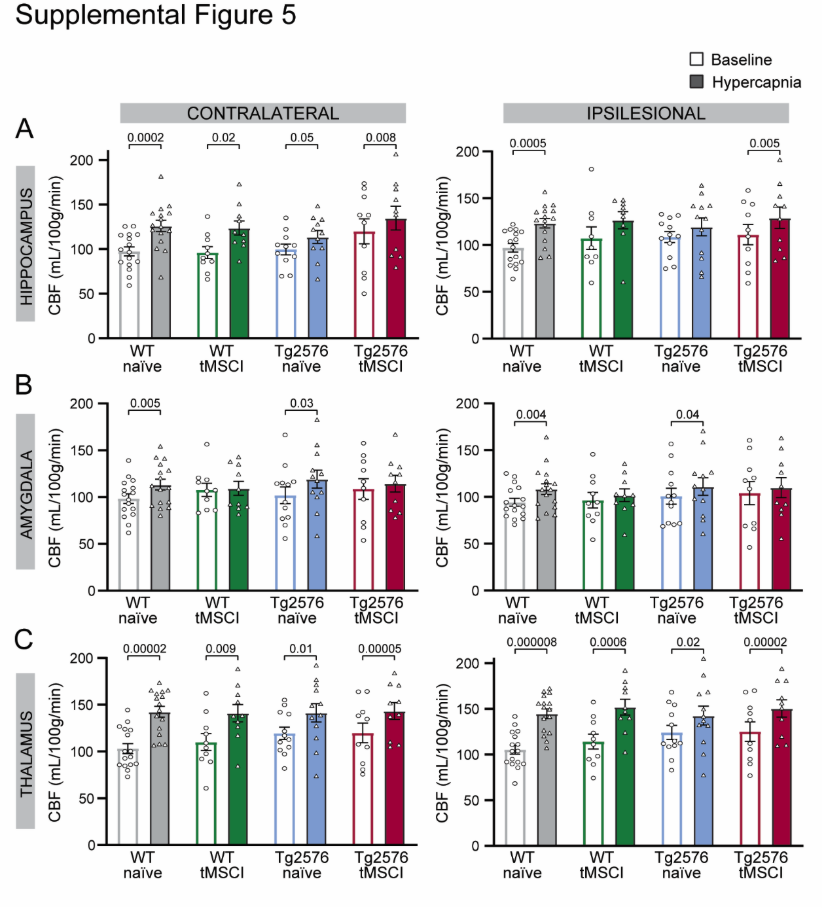

### Supplemental Figure 5. Cerebrovascular reactivity is impaired in the hippocampus and amygdala but not the thalamus at 8 mo post-tMSCI.

**(A-C)** Hypercapnic hyperemia displayed as raw CBF values at baseline (circles) and following hypercapnia (triangles) for contralateral and ipsilateral hippocampus **(A)**, amygdala **(B)**, and thalamus **(C)**. Data were compared within each experimental group using paired t-tests.

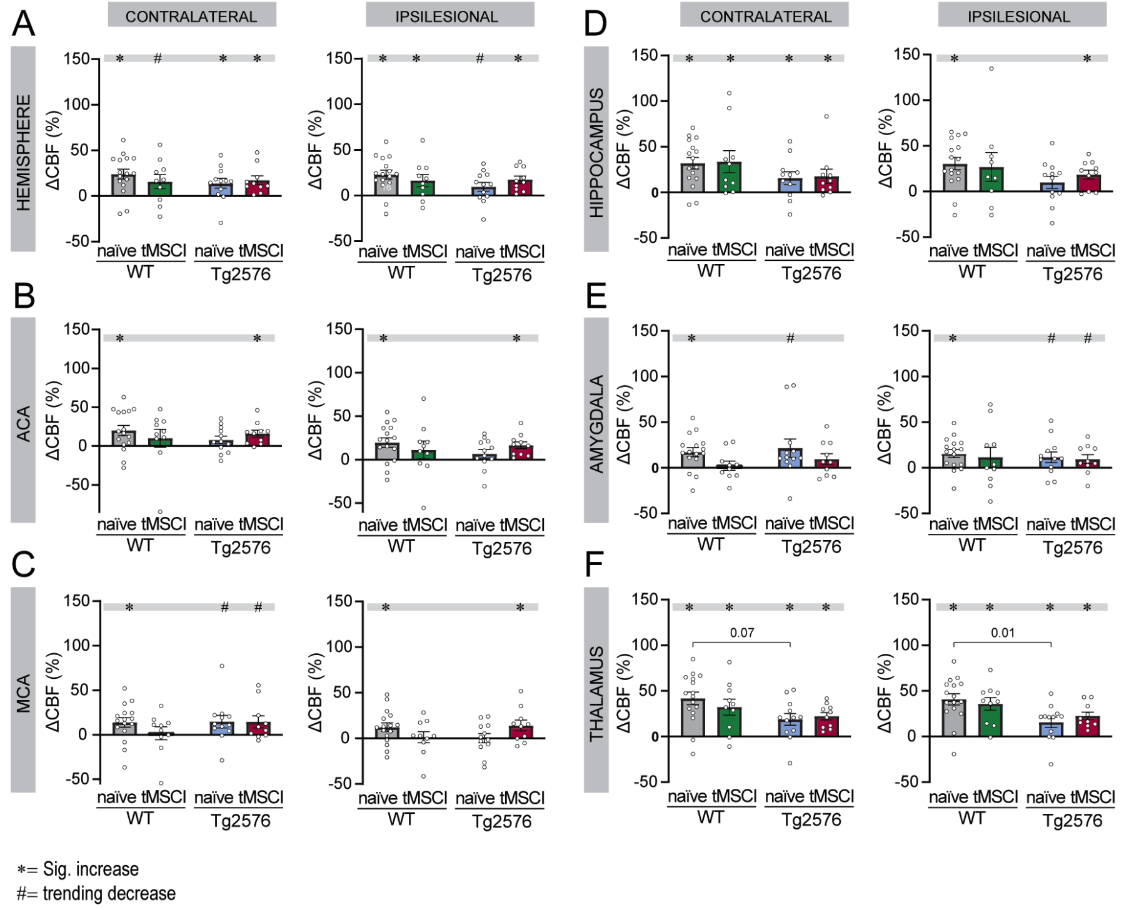

### Supplemental Figure 6. Aβ and stroke differentially impair cerebrovascular reactivity by region at 8 mo post-tMSCI.

**(A-F)** Hypercapnic hyperemia displayed as a percentage of baseline CBF for contralateral and ipsilateral hemisphere **(A)**, ACA **(B)**, MCA **(C)**, hippocampus **(D)**, amygdala **(E)**, and thalamus **(F)**. Data were compared within each experimental group using one-way ANOVAs with Tukey’s multiple comparisons. Symbols in gray bars depict results of one-sample t-tests comparing data to no change in CBF. * indicates significant increase (p<0.05), # indicates a trending increase (p<0.1).

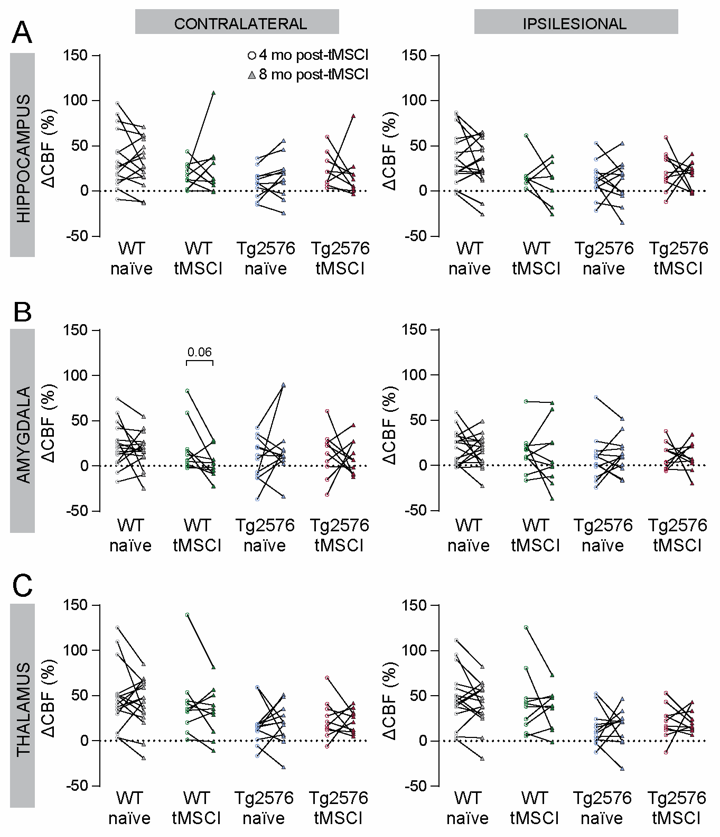

### Supplemental Figure 7. Longitudinal analysis shows that cerebrovascular reactivity is not significantly altered between 4 to 8 mo post-tMSCI in hippocampus, amygdala, and thalamus.

**(A-C)** Hypercapnic hyperemia displayed as a percentage of baseline CBF at 10 mo (circles) and 14 mo (triangles) for contralateral and ipsilateral hippocampus **(A)**, amygdala **(B)**, and thalamus **(C)**. Data were compared within each experimental group using multiple paired t-tests.

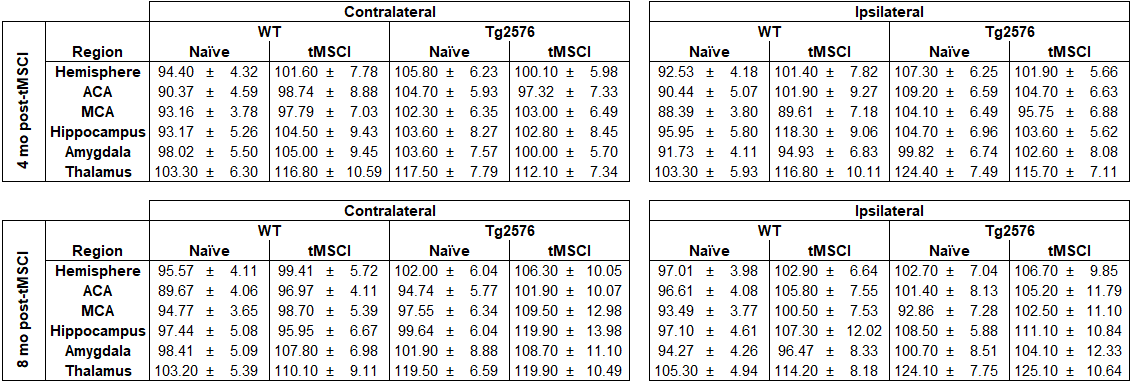

### Supplemental Table 1. Resting levels of CBF at 4 and 8 mo post-tMSCI.

Average resting CBF measurement ± SEM (mL/100 g/min) by brain region from each hemisphere of each group at both timepoints. No differences were found in regional CBF between groups at either timepoint (one-way ANOVA).
